# Chromosome-level genome assembly of the swallowtail butterfly *Parides eurimedes mylotes* is a valuable resource for studying wing coloration

**DOI:** 10.64898/2026.05.27.728091

**Authors:** Praveen Kumar Jaya Balaji, Tim Davalan, Pamela Nicholson, Carmen Rojas Uglade, Laurent Falquet, Viola Vogler-Neuling

## Abstract

High-quality chromosome-scale assemblies are scarce in *Papilionidae*. This limits comparative genomics to model species, *Lepidoptera, Bombyx mori*. Here, we present a phased, chromosome-level genome assembly of *Parides eurimedes mylotes*. We generated this assembly using 125⨯ PacBio HiFi sequencing and assembled it with hifiasm. The final haplotype assemblies (Hap1 and Hap2) span 274 Mb and 270 Mb, respectively. These assemblies are organized into 31 near-telo-mere-to-telomere chromosomes, with scaffold N50 values of 9.72 Mb and 9.22 Mb, respectively. BUSCO analysis revealed assembly completeness of 96.6 % and 96.4 % for Hap1 and Hap2, respectively. Repeats annotation identified 18-19 % repetitive content, with Helitron elements being the dominant class of transposable elements. We identified the W and Z sex chromosomes and completely assembled the mitochondrial genome. Compared to the previously available *Parides photinus* draft assembly, our genome exhibits an 11,000-fold reduction in scaffold fragmentation and nearly complete gene assembly. This assembly provides a robust genomic reference for functional, evolutionary, and multi-omics investigation in *Papilionidae*.

In addition to serving as a high-quality genomic reference for *Papilionidae*, this assembly is essential for linking the genetic architecture of butterfly wings to the hierarchical nanostructures underlying structural coloration. By identifying the genes and regulatory networks involved in scale morphogenesis, we can correlate the butterfly’s genotype with its photonic function. This insight into the evolutionary origin of the biological photonic systems informs the design of biomimetic, structurally colored materials.

## Background & Summary

Unlike conventional pigments, which rely on light absorption, structural color arises from the interference, diffraction, and scattering of light within nanostructures. This color-producing strategy is widely used by organisms in nature for functions such as communication, mate selection, camouflage, and survival^1^.

In these systems, optical effects are governed by periodically ordered nanoscale structures that form during insect development. Butterflies, for example, have diverse nanostructures, including thin films, ridges, smooth multilayers, and 3D gyroid structures^2^. While many of these structures are well characterized, the molecular building blocks of these architectures, including lipids and proteins, as well as the genes and regulatory networks responsible for these nanostructures, remain largely unidentified. This leaves a significant genotype-phenotype gap in the field of structural colors. This gap hinders the biomimetic replication of biological photonic crystals.

Butterfly species within the genus *Parides* (cattleheart butterflies) are an ideal model system for addressing this gap, because they exhibit striking diversity in photonic nanostructures on their wing scales within the same phylogenetic lineage. Reported scale morphologies within this genus include smooth multilayers, ridges, and 3D gyroids^3^. Importantly, these nanostructures co-exist with pigmentary elements, suggesting coupled optical mechanisms that are poorly understood at the molecular level. This structural and optical diversity, combined with their close evolutionary relationships, makes *Parides* a promising model for elucidating the genetic and molecular processes underlying distinct photonic structures and their optical functions.

Although advances in long-read sequencing technologies have rapidly advanced the field of lepidopteran genomics, many non-model butterfly genomes remain fragmented and incomplete. Current genomic resources for butterflies, including those for the monarch (*Danaus plexippus*)^4^, *Bicyclus anynana*^5^, and *Heliconius melpomene*^6^, have primarily advanced our understanding of pigmentation and wing patterning. However, these resources focus largely on pigment-based coloration rather than structural coloration. Historically, genomic resources within *Lepidoptera* have focused heavily on the silkworm, *Bombyx mori*^7–9^. Additionally, the *Bombyx mori* genome contains a high percentage of repetitive sequences (> 30 %)^10,11^, which limits comparative and lineage-specific analyses across *Papilionidae*.

Although *Bombyx mori* is a foundational model, its deep evolutionary divergence^12,13^ makes it an insufficient reference for studying lineage-specific traits, such as the photonic nanostructures of butterflies. Using distant references, such as *Bombyx mori*, in proteomics studies of non-model organisms can yield inaccurate results due to evolutionary divergence, sequence variability, and the absence of lineage-specific structural proteins. This can result in misidentifications and the loss of information about the exact proteins responsible for nanostructure formation, severely hindering mechanistic insight.

The genus *Parides* is scientifically and evolutionarily significant within the *Papilionidae* family. However, genomic resources are sparse and fragmented. The previously assembled *Parides photinus* genome (*Genbank AC:GCA_018247975*.*1*) consisted of more than 300,000 scaffolds, only 73 % of which were complete^14,15^, limiting chromosomal inference.

For this study, *Parides eurimedes mylotes* was selected based on biological relevance and experimental feasibility. This species has a well-defined lumen with a smooth multilayered structure, making it an ideal candidate for studying the genetic basis of structural coloration in the *Papilionidae*^3^. Unlike many rare, endangered, and protected species with interesting optical features, *Parides eurimedes mylotes* is not endangered and is accessible to authorized breeders. This allows for statistically robust downstream analyses, including proteomics and lipidomics, which require biological replicates to determine the structure-function relationship with high confidence.

Recent advances in high fidelity (HiFi) long-read sequencing, combined with graph-based assembly algorithms, now enable phased, chromosome-scale assemblies with improved resolution and haplotype separation. These assemblies provide greater gene completeness and an accurate representation of transposable elements within the gene sequence and the overall structural architecture.

Here, we present a chromosome-level, phased genome assembly of *P. eurimedes mylotes* using deep PacBio HiFi sequencing. We evaluate the assembly’s continuity, BUSCO completeness, repeat architecture, sex chromosome identification, and improvements compared to previously available *Parides* genomic resources. Our goal is to establish a high-quality genome reference for the *Papilionidae* family that will enable future evolutionary and functional studies. This will help us identify the genes responsible for photonic structures. Additionally, these data can be used to annotate protein building blocks and establish a phylogenetic tree. By linking genotype, structural, and optical function, our work provides a foundation for the biomimetic fabrication of these photonic structures using biological building blocks – a goal that remains elusive for the photonic and material science communities.

Beyond providing a foundation for studying photonic nanostructures, this genomic resource opens multiple avenues for interdisciplinary research. It allows us to investigate genetic pathways that govern scale morphogenesis, cuticular organization, and hierarchical structural assembly at the nano- and microscale. These pathways govern functional properties, such as hydrophobicity, and the development of lightweight, mechanically robust wing architecture. Phased, chromosome-level information facilitates the study of regulatory evolution, structural variation, and sex-linked traits. It also supports the accurate annotation of protein building blocks involved.

## Methods

### Sample collection and sequencing

On April 28, 2025, butterfly pupae were purchased from The Entomologist Ltd., located at Middleton Common Farm in East Sussex. The pupae were purchased under the name *Parides arcas*. The company is a member of the International Association of Butterfly Exhibitors and Suppliers (IABES), which protects wild butterflies and their habitats by promoting sustainable exhibits and supporting conservation, research, and public education. Carmen Rojas Ugalde, from the Universidad de Costa Rica, analyzed the pupae and identified them as *Parides eurimedes mylotes* (H. W. Bates, 1861). The pupae were raised in a humidity- and temperature-controlled terrarium until they eclosed. Figure 1A and D show the female and male imagoes of *Parides eurimedes mylotes* butterflies. The female imago was frozen at –20 °C. After at least three days at this temperature, the butterfly was dissected into four parts: the body (**Figure 1G**), the forewings (FW) (**Figure 1H**), the yellow-colored patches (**Figure 1I**), and the hindwings (HW) (**Figure 1K**). These parts were submitted for genome sequencing.

**Figure 1.**
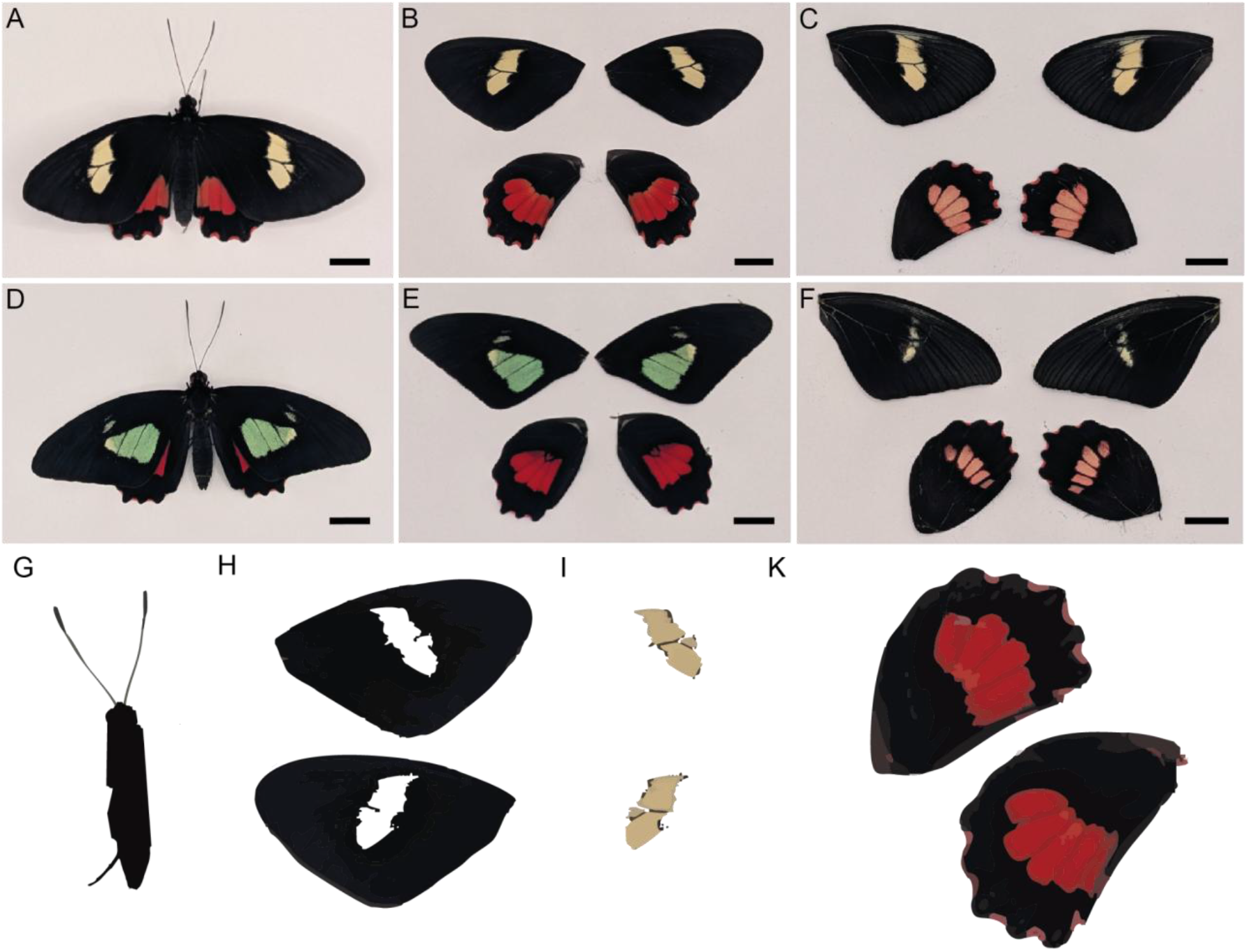
**A** Specimen picture of the female *Parides eurimedes mylotes* imago used for genome sequencing. **B** Forewings of female; **C** Hindwings of female. **D** Specimen picture of male imago. **E** Forewings of male; and **F** Hindwings of male. The female image was dissected into the following parts: **G** Body, **H** forewings, **I** yellow patches, and **K** hindwings. Scale bars: all 1 cm.

We extracted high-molecular-weight (HMW) genomic (g)DNA using the 10⨯ genomics sample preparation demonstrated protocol, “10⨯ Genomics Sample Preparation Demonstrated Protocol DNA Extraction from Single Insects” (10⨯ Genomics, CG000145, Rev A)^16,17^. Prior to SMRTbell library preparation, we assessed the quantity, quality, and purity of the DNA using a Qubit 4.0 fluorometer (Qubit dsDNA HS or BR Assay kit; Q32851/Q32850, Thermo Fisher Scientific), an Advanced Analytical FEMTO Pulse instrument (Genomic DNA 165 kb Kit; FP-1002-0275, Agilent), and a Denovix DS-11 UV-Vis spectrophotometer, respectively. SMRTbell libraries were prepared according to the PacBio guideline, “Preparing whole genome and metagenome libraries using SMRTbell prep kit 3.0” (PacBio,102-166-600 REV04 APR 2024). Two extracted samples (FW and HW) were selected for library preparation (**Figure 1H** and **I**). Briefly, sheared gDNA was concentrated and cleaned using 1⨯ SMRTbell clean-up beads. The samples were quantified and confirmed to be in the 13–20 kb range using a Qubit 4.0 fluorometer (Qubit dsDNA HS Assay kit; Q32851, Thermo Fisher Scientific) and an Advanced Analytical FEMTO Pulse instrument (Genomic DNA 165 kb Kit; FP-1002-0275, Agilent), respectively. The rest of the aforementioned procedure was followed, including end-repair and A-tailing, ligation of barcoded overhang adapters, library purification using AMPure PB beads to remove DNA fragments below 5 kb. A nuclease treatment was also performed. Final libraries were checked for quantity and quality using a Qubit 4.0 fluorometer (Qubit dsDNA HS Assay kit; Q32851, Thermo Fisher Scientific) and an Advanced Analytical FEMTO Pulse instrument (Genomic DNA 165 kb Kit; FP-1002-0275, Agilent). The two libraries were equimolarly pooled and rechecked for quantity and size, as outlined above. Instructions in SMRT Link Sample Setup were followed to prepare the SMRTbell library for sequencing (PacBio SMRT Link v25.2). The PacBio standard sequencing primer was annealed to the SMRTbell libraries from a Revio SPRQ polymerase kit (PacBio, PN 103-496-900). Then, Revio DNA polymerase was bound, and the polymerase-bound complex was purified using cleanup beads (PN 102-158-300, PacBio). Finally, the Revio sequencing control DNA was diluted and added to the complex before pipetting onto the thawed Revio SPRQ sequencing plate (PacBio, PN 102-326-552). The Revio deck was set up as directed by the SMRTLink software. This included laying out tips, sequencing plates, and Revio SMRT Cell trays. Each tray contained four 25 M SMRT cells (PacBio, PN 102-202-200). The libraries were loaded at an on-plate concentration of 300 pM using adaptive loading. SMRT sequencing was performed on the Revio for a 30-hour movie time. All steps, from gDNA extraction to post-run data utilities such as demultiplexing, were performed at the Next Generation Sequencing Platform at the University of Bern, Switzerland.

### Genome assembly

First, we performed quality control (QC) of the reads using FASTQC^18^. Then, we cleaned the reads with fastp^19^. Next, we concentrated the cleaned reads from the four wing samples (two forewings and two hindwings). We performed genome assembly on these reads with hifiasm^20^ with the default parameters as well as the “--telo-m TTAGG” parameter to account for the canonical telomeric sequences of Lepidoptera. Two haplotigs, Hap1 and Hap2, were obtained.

We validated the assembled haplotigs using the following tools: BUSCO^21^, MERQURY^22^, GenomeScope2^23^, Tidk^24^, EDTA^25^, chromsym^26^, nucmer^27^, DOT^28^.

The assembly quality was excellent, with 27 T2T chromosomes identified as well as five fragmented chromosomes that could be joined using Seqkit^29^. A total of 2*n* = 60 chromosomes, plus the Z and W sexual chromosomes, were reconstructed (as expected for a female). We identified and circularized the mitochondrial genome by comparing remaining contigs with the mitochondrial genome of *Parides photinus* (ACnum: LS974638) ^5^.

### Genome annotation

We annotated the two haplotigs’ chromosomes independently using the EGAPx pipeline from NCBI (https://github.com/ncbi/egapx; version 0.5.0) and RNAseq data from *Ornithoptera alexandrae* as a proxy for *P. eurimedes mylotes* (PRJNA938052; SRR23685258_1; SRR23685258_2). Our analysis revealed 11,402 CDSs for Hap1, 11,156 CDSs for Hap2, as well as six non-coding RNAs for both haplotypes. We annotated the proteins using InterProScan v5.77-108.0.

## Results

### Genome sequencing and assembly

We sequenced HMW genomic DNA from *P. eurimedes mylotes* with approximately 125× coverage using PacBio HiFi chemistry. *De novo* assembly using hifiasm generated phased primary (Hap1) and alternate (Hap2) assemblies. The final Hap1 assembly spans 274 Mb, with an N50 value of 9.72 Mb, while the Hap2 assembly spans 270 Mb, with an N50 value of 9.22 Mb. Both assemblies resolved 31 large scaffolds corresponding to chromosome-scale sequences. These metrics represent a substantial improvement in contiguity compared to previously published *Parides* assemblies.

### Assembly quality and completeness

We assessed assembly completeness using BUSCO v6.0.0 with the Lepidoptera_odb12 dataset. Hap1 achieved 96.6 % completeness with only 3.3 % of the genes missing. Hap2 achieved 96.4 % completeness. These results are a significant improvement over those of *Parides photinus* (Genbank AC:GCA_018247975.1)^30^, which had 74.8 % complete BUSCOs and 12.3 % missing genes. The reduction in missing orthologs indicates improved gene-space representation and reduced fragmentation^31,32^. GenomeScope^33^-based *k*-mer analysis estimated a smaller genome size than that of the assembled haplotypes (**Figure 2**). The estimated sizes were ~238 Mb for Hap1 and 274 Mb for Hap2. This discrepancy may be due to ~19 % repeats.

**Figure 2.**
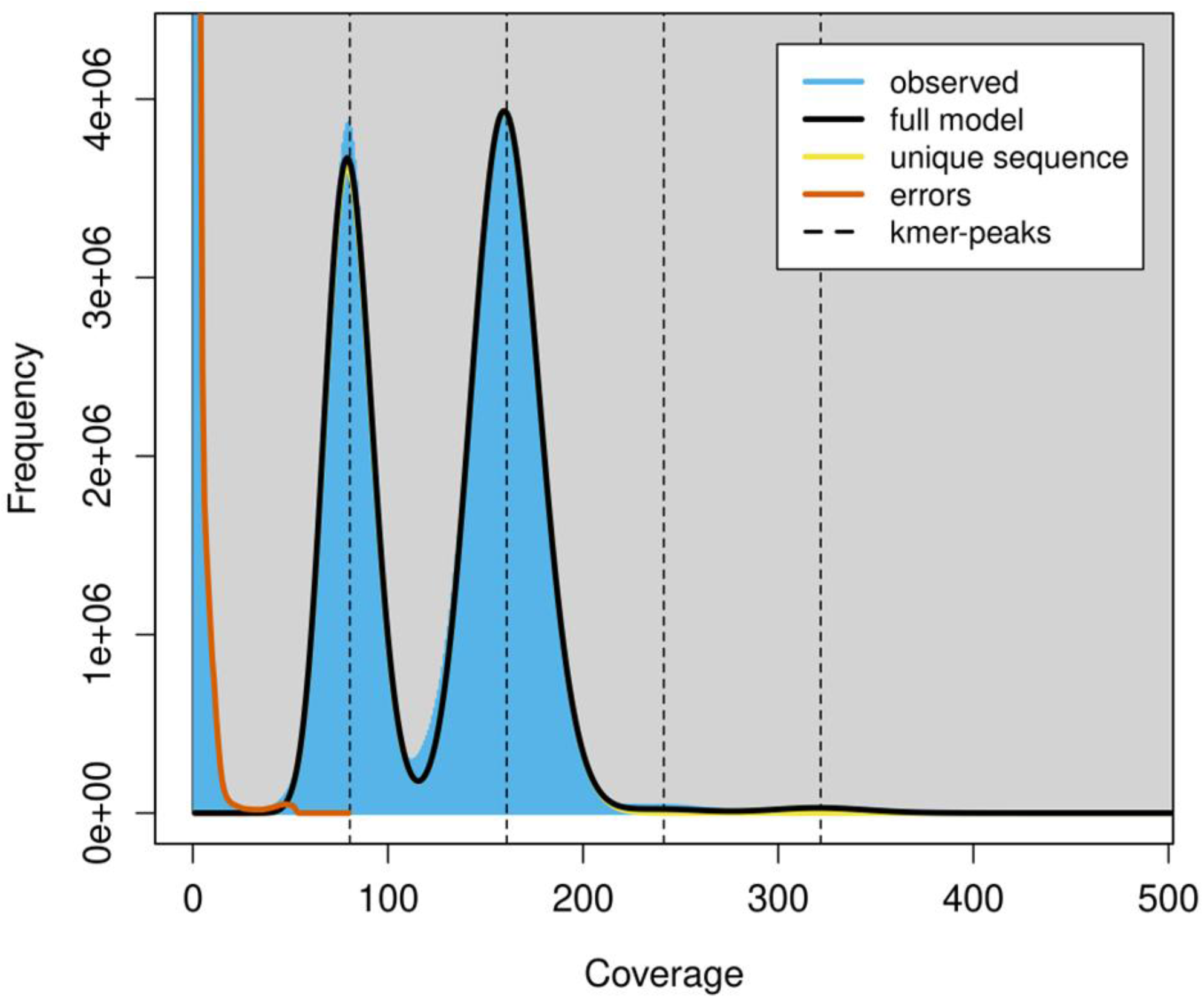
*k*-mer frequency distribution used to estimate genome size and heterozygosity estimation with GenomeScope 2.0 and a *k*-value of 21.

The GenomeScope plot compares *k*-mer abundances (blue) with a mathematical model (black line) used to estimate the genome’s overall characteristics. The profile exhibits a clear bimodal distribution represented by two blue peaks, indicative of a diploid organism. The first peak, known as the half-coverage peak, represents heterozygous *k*-mers. The second, taller peak represents homozygous regions of the genome. Based on this profile, the estimated haploid genome size is approximately 238 Mb with a high sequence uniqueness of ~94 % and minimal duplication (~0.85 %). The genome sequence’s heterozygosity is around 1.33 %, indicating moderate genetic variation. Most importantly, the sequencing error rate is very low at ~0.078 %, as shown by the orange line. The observed sequence coverage also fits the model well.

### Repeat landscape characterization

Repeat annotation was performed using EDTA. Approximately 19 % of the *P. eurimedes mylotes* genome consists of repetitive elements. Helitrons were identified as the dominant transposable element class, accounting for nearly 8 % of the genome. Other detected elements include long interspersed nuclear elements (LINEs), Long terminal repeat (LTR) retrotransposons, and DNA transposons^34^ (Suppl. Table 1). Although the proportion of repeats is lower than in some reported Lepidopteran genomes^34–36^, it falls within the observed range of *Papilionidae*.

### Chromosome architecture and sex chromosome identification

The assembly produced 31 chromosome-scale scaffolds consistent with known Lepidopteran karyotypes. A comparative analysis of coverage and gene content revealed that two of the scaffolds correspond to the W and Z sexual chromosomes. Synteny plots comparing the two haplotigs to *Bombyx mori* chromosomes show excellent synteny (Figures 3 and 4).

**Figure 3.**
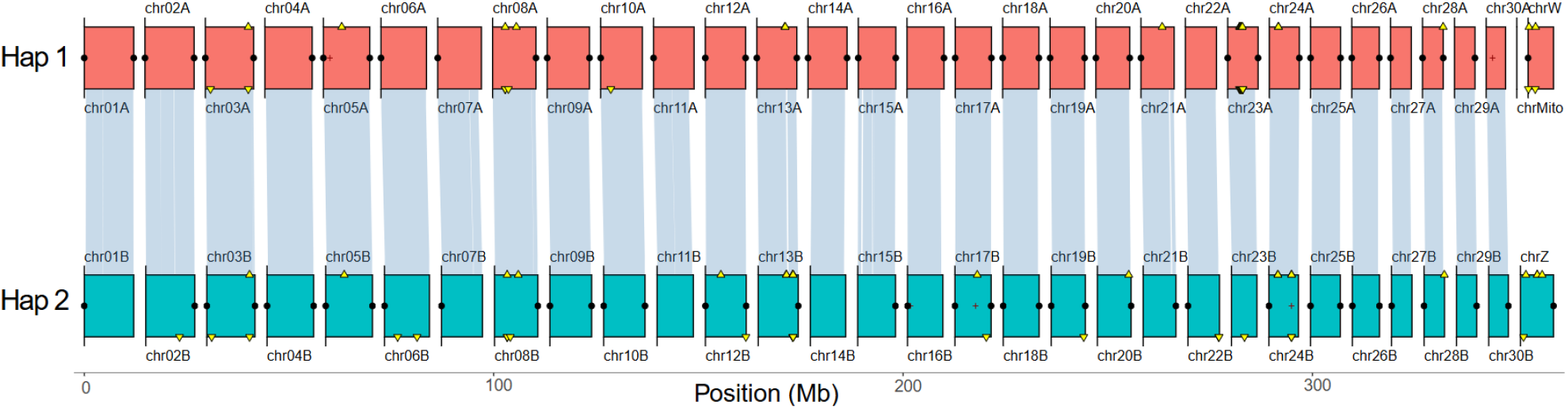
Synteny plot of the two *Parides eurimedes mylotes* haplotigs. Haplotig 1 (red) and 2 (turquoise) were examined for synteny. Red crosses along chromosomes represent assembly gaps, and black dots at the ends indicate telomere sequences. Yellow triangles indicate duplicated BUSCO genes in haplotigs 1 and 2. Blue links between assemblies represent collinear regions, while red links represent genome inversions.

**Figure 4.**
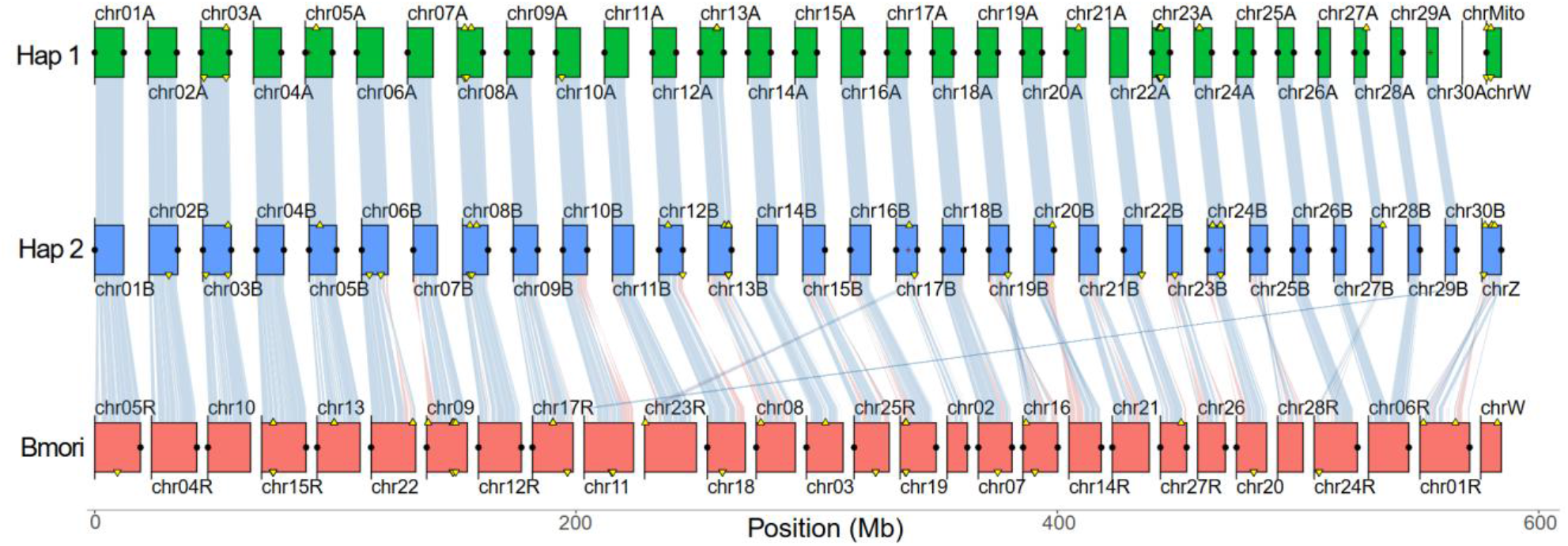
Synteny plot of the *Bombyx mori* chromosome (ASM2736675v1)^37^. Haplotigs 1 (green) and 2 (blue), generated in this study, were tested for synteny against the *Bombyx mori* reference genome (red). Chromosomes with names ending in “R” are inverted. The genome assembly shows high synteny at the chromosome level with *Bombyx mori* and a high degree of conservation across both haplotypes. Minor localized inversions and rearrangements reflect evolutionary divergence rather than assembly error.

### Mitochondrial genome assembly

We assembled a complete mitochondrial genome using HiFi reads. The gene content and organization were consistent with the canonical Lepidopteran mitochondrial architecture as annotated using Proksee.ca^38^ (**Figure 5**).

**Figure 5.**
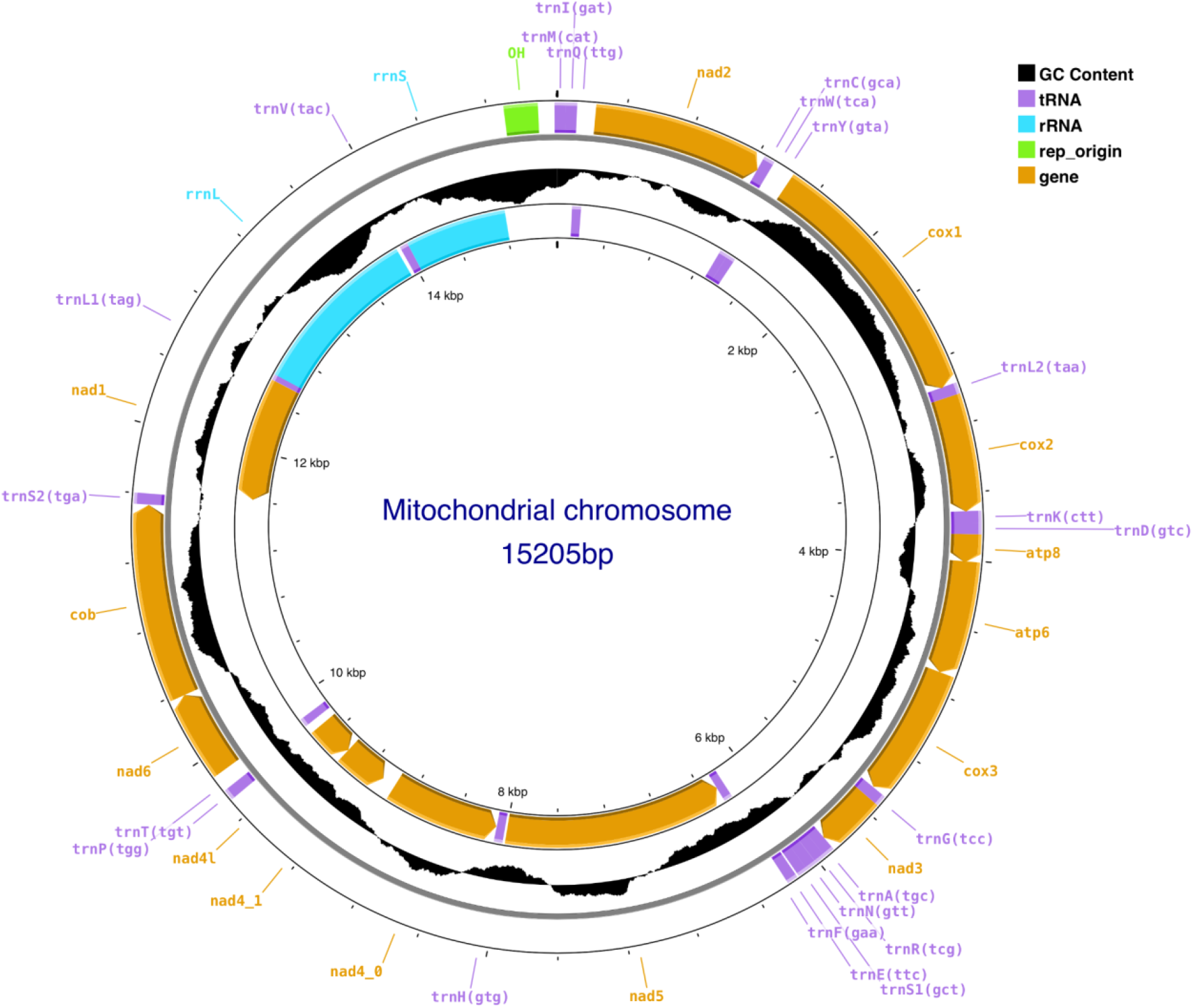
Annotated mitochondrial genome of *Parides eurimedes mylotes* using Proksee.ca

### Comparative assessment with *Parides photinus*

The previously published *P. photinus* assembly comprised 338,000 highly fragmented scaffolds and showed reduced BUSCO completeness (~74 %). In contrast, our study of *P. eurimedes mylotes* revealed a reduction in the number of scaffolds by about four orders of magnitude (from ~3.4 × 10^5^ to ~3.2 × 10^1^), an increase in BUSCO completeness of 22 %, and a substantial reduction in missing orthologs. Additionally, we observed chromosome-scale contiguity and enhanced repeat resolution (**Figure 6**).

**Figure 6.**
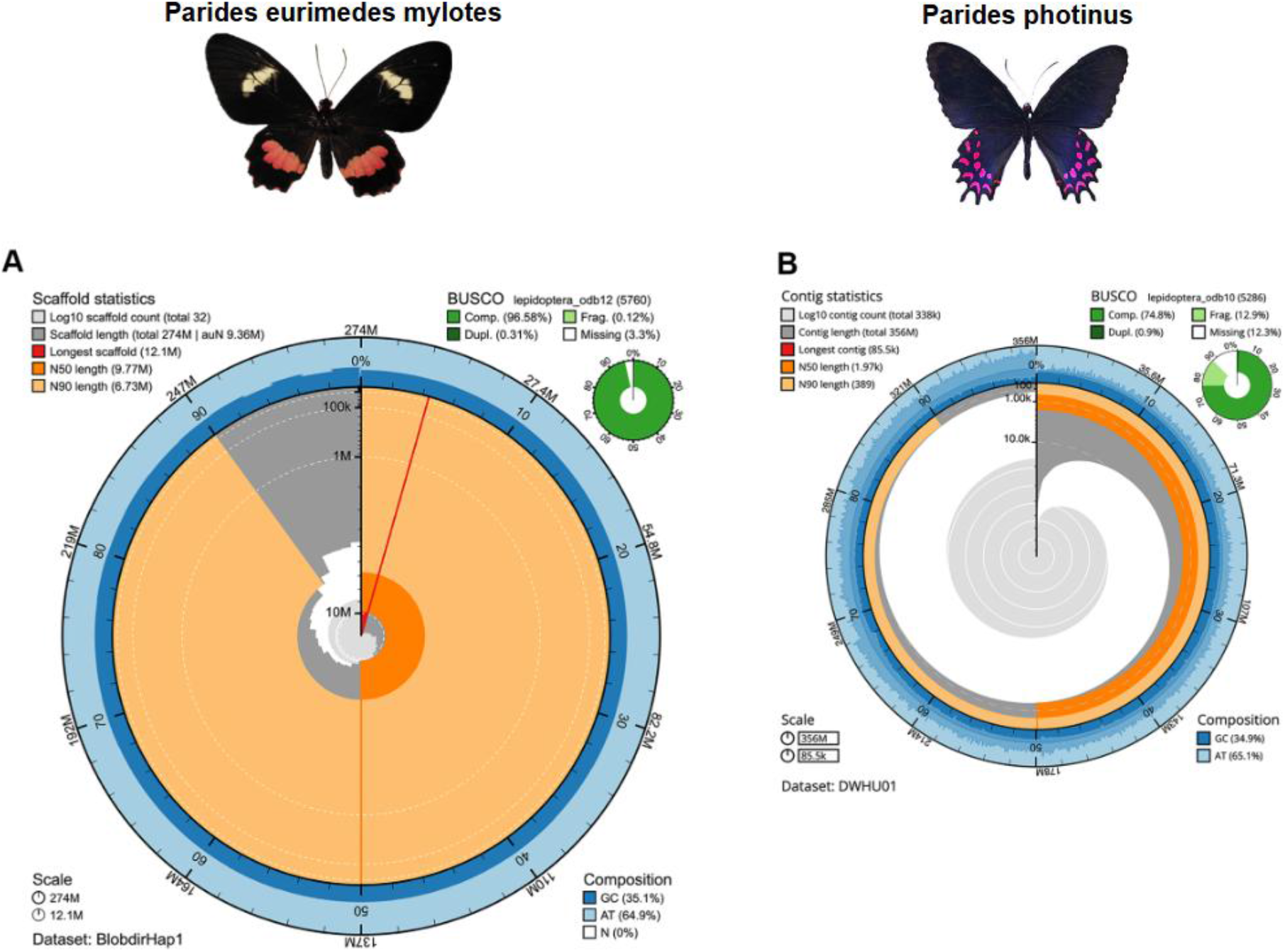
Snail plots: **A** *Parides eurimedes mylotes* Hap1 and **B** *Parides photinus*. Assemblies were obtained from https://blobtoolkit.genomehubs.org/view/Parides/dataset/DWHU01/snail. Hap2 is not shown but is nearly identical to Hap1 (see Suppl. Figure 1).

### Potential genes involved in wing development and coloration identified

A set of genes associated with pigmentation, cuticle formation, cytoskeletal organization, and membrane trafficking was identified in the *Parides eurimedes mylotes* genome assembly. Two annotation pipelines, EGAPx and SANSPANZ, were used to identify these genes across the Hap1 and Hap2 haplotypes (**Table 1**). The two pipelines’ annotations tend to agree. Notably, the transcription factor optix^39^ was detected in the SANSPANZ annotation and appears once in each haplotype. However, the EGAPx annotation identified it as the homeobox six6 gene. The morphogen gene^40^ was present in two copies in both EGAPx haplotypes and in one copy in the SANSPANZ annotation. Several genes associated with cuticle formation were identified, including chitin synthase^41^, which was represented by two copies per haplotype in EGAPx and by one copy in SANSPANZ. The CPAP/peritrophin genes^42^ were also identified and represented by a single copy in both pipelines. Cytoskeletal genes such as F-actin^43^ (one copy per subunit and per haplotype in EGAPx and in SANSPANZ), formin^44^, and cofilin (one copy in both annotations) were also identified. Rab5, Rab7, and Rab11^45^ membrane trafficking genes were identified in both annotation sets, with similar copy numbers across haplotypes. Several genes involved in pigmentation pathways were also present, including vermilion, kynurenine pathway, white, yellow, ebony, melanization protease, and tyrosine metabolism genes. DDC laccase^46–49^ was present at varying copy numbers in the two annotation pipelines, though it was consistent across the two haplotypes. Interestingly, the ebony and tyrosine hydroxylase genes were present as a single copy only on Hap1 and absent from Hap2. The number of yellow and white genes was unclear. Overall, the surveyed gene set was detected in the genome assembly, and little difference in gene copy counts was observed between the two annotation approaches.

**Table 1.**
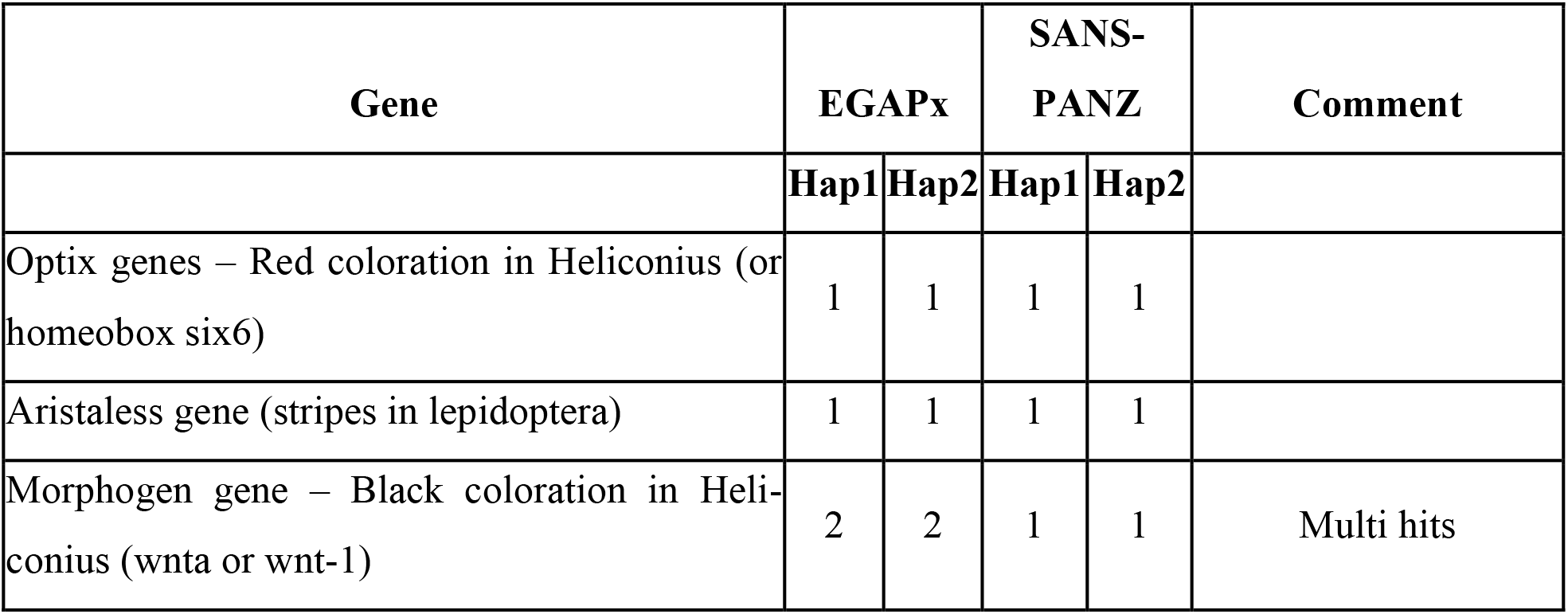
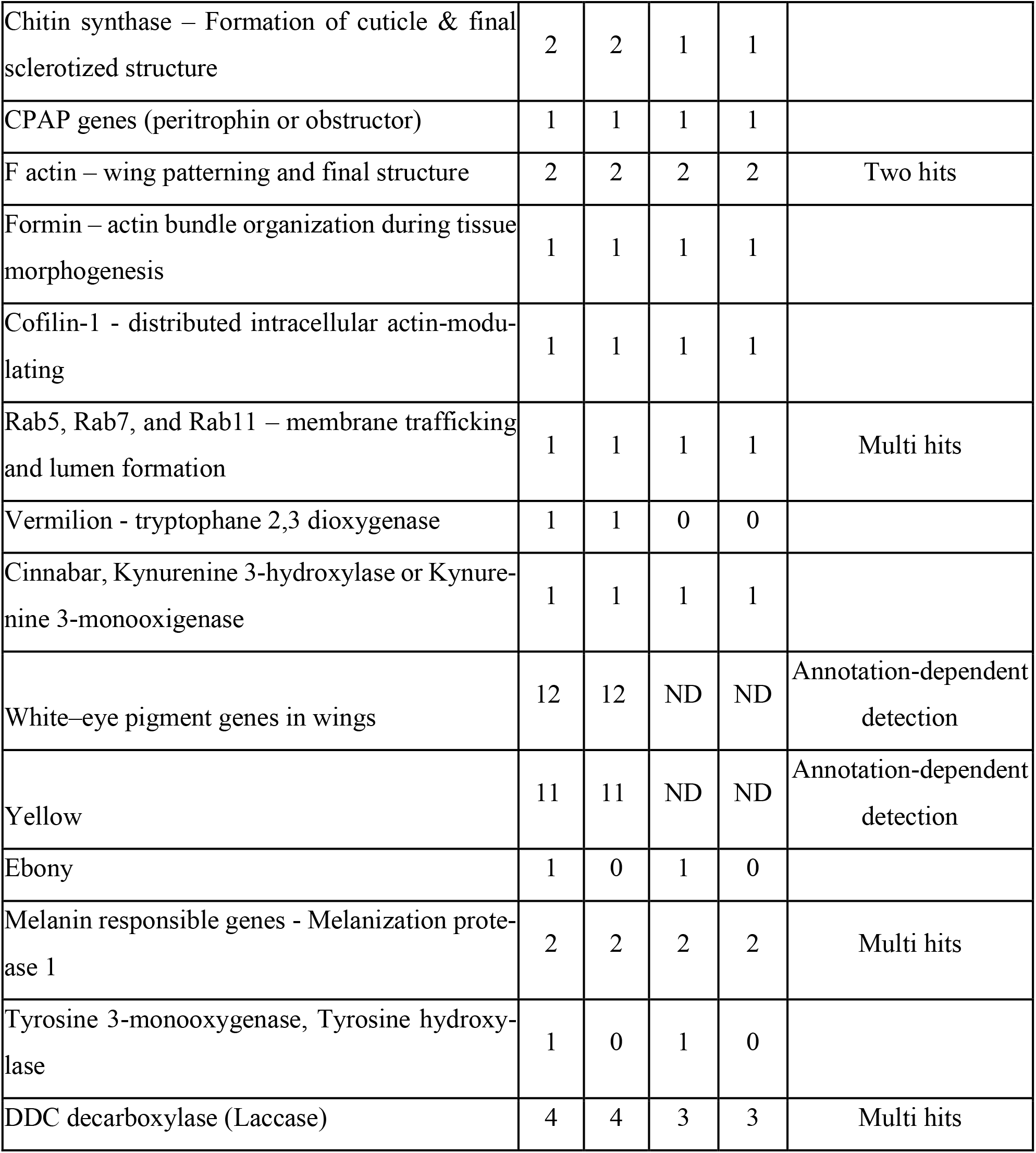
Detection of candidate genes associated with coloration, cuticle formation, and cellular organization across both haplotypes using two independent annotation pipelines (EGAPx and SANSPANZ). ND indicates not determined.

| Gene | EGAPx |  | SANS-PANZ |  | Comment |
| --- | --- | --- | --- | --- | --- |
|  | Hap1 | Hap2 | Hap1 | Hap2 |  |
| Optix genes – Red coloration in Heliconius (or homeobox six6) | 1 | 1 | 1 | 1 |  |
| Aristaless gene (stripes in lepidoptera) | 1 | 1 | 1 | 1 |  |
| Morphogen gene – Black coloration in Heliconius (wnta or wnt-1) | 2 | 2 | 1 | 1 | Multi hits |
| Chitin synthase – Formation of cuticle & final sclerotized structure | 2 | 2 | 1 | 1 |  |
| CPAP genes (peritrophin or obstructor) | 1 | 1 | 1 | 1 |  |
| F actin – wing patterning and final structure | 2 | 2 | 2 | 2 | Two hits |
| Formin – actin bundle organization during tissue morphogenesis | 1 | 1 | 1 | 1 |  |
| Cofilin-1 - distributed intracellular actin-modulating | 1 | 1 | 1 | 1 |  |
| Rab5, Rab7, and Rab11 – membrane trafficking and lumen formation | 1 | 1 | 1 | 1 | Multi hits |
| Vermilion - tryptophane 2,3 dioxygenase | 1 | 1 | 0 | 0 |  |
| Cinnabar, Kynurenine 3-hydroxylase or Kynurenine 3-monooxygenase | 1 | 1 | 1 | 1 |  |
| White-eye pigment genes in wings | 12 | 12 | ND | ND | Annotation-dependent detection |
| Yellow | 11 | 11 | ND | ND | Annotation-dependent detection |
| Ebony | 1 | 0 | 1 | 0 |  |
| Melanin responsible genes - Melanization protease 1 | 2 | 2 | 2 | 2 | Multi hits |
| Tyrosine 3-monooxygenase, Tyrosine hydroxylase | 1 | 0 | 1 | 0 |  |
| DDC decarboxylase (Laccase) | 4 | 4 | 3 | 3 | Multi hits |

The identification of these genes confirms that the assembled genome contains key pathways associated with pigmentation, cuticle formation, and cytoskeletal organization. Differences in predicted gene copy numbers between EGAPx and SANSPANZ likely reflect differences in gene model prediction between the two annotation pipelines, as the number of genes remains consistent across the haplotypes.

Table 1 summarizes the gene copy numbers identified in Hap1 and Hap2 using both annotation methods. Developmental and pigmentation-related genes, such as *optix, wntA*, and melanin pathway genes, were consistently detected in both haplotypes, supporting the completeness of the phased assembly. However, differences in gene counts between annotation methods reflect variation in prediction sensitivity, as some genes show multiple hits or duplicated annotations. White-eye pigment genes and yellow genes were detected in both haplotypes using EGAPx but not the SANSPANZ annotation pipeline. These discrepancies are a known limitation of automated annotation approaches and do not indicate the true absence of a gene from the genome. Therefore, further validation is required to confirm the presence and the copy number of these genes.

## Discussion

We present a high-quality, chromosome-scale genome assembly of *Parides eurimedes mylotes* that is over 96 % complete, surpassing all previously published *Parides* assemblies. Using ~125× PacBio HiFi data and hifiasm, we generated phased Hap1 and Hap2 assemblies with N50 scaffold lengths of 9.72 and 9.22 Mb, respectively. These assemblies resolved 31 chromosome-scale scaffolds, including the Z and W sex chromosomes. The Hap1 assembly achieved 96.6 % BUSCO completeness, demonstrating near-complete gene space representation and substantially fewer missing orthologs than the 75 %-complete *P. photinus* assembly.

This increase in gene completeness has critical functional consequences. Fragmented and incomplete assemblies obscure gene families, collapse repetitive regions, and disrupt syntenic relationships. These issues hinder the accurate identification of structural variants, regulatory elements, and gene duplications. The nearly complete, phased assembly presented in this study enables reliable detection of these features. It also facilitates chromosome-level comparative genomics with *Papilionidae* and provides the necessary resolution to correlate genetic architecture with the formation of photonic nanostructures.

The genome spans ~274 Mb (Hap1), exceeding the estimated 238 Mb from GenomeScope using *k*-mers. This discrepancy is likely due to repeats, as annotation shows that approximately 19 % of the genome is repetitive, dominated by helitrons (~8 %).

Our assembly reconstructs 31 T2T chromosomes and a mitochondrial genome, providing a framework for functional and comparative genomics. Compared with the highly fragmented published assembly of *P. photinus*, our work demonstrates a 10^4^ reduction in scaffold count, 22 % higher BUSCO completeness, and improved repeat resolution.

This genome study establishes a foundational reference for *Papilionidae* genomics, moving beyond the traditional *Bombyx mori* reference. Furthermore, this assembly is a critical step toward linking genotype to phenotype in structural-colored butterflies. As the first near-complete gene assembly of a structurally colored butterfly, it allows us to examine the genetic basis of photonic wing structures and identify lineage-specific genes involved in scale morphogenesis. It also allows us to integrate transcriptomics to map the developmental dynamics of nanostructures and to conduct comparative analyses to determine whether the genetic architectures underlying photonic structures on butterfly wings are conserved or divergent.

Subsequent proteomic analyses, facilitated by this assembly, will address a key bottleneck in understanding the nanostructure formation by identifying the molecular building blocks of nanostructures. These genomic and proteomic insights can inform biomimetic design by translating naturally evolving structural color systems into engineered photonic materials. This approach bridges the gap between evolutionary biology and photonic material engineering.

## Data Records

The raw genome reads are deposited at ENA under project PRJEB108058.

## Acknowledgment

The authors acknowledge the funding provided to Praveen Kumar Jaya Balaji through the industrial PhD program “Talents for Biobased Economies (T4BBI),” a Marie Skłodowska-Curie COFUND project under the EU Horizon 2020 Research and Innovation (grant number 101034323). BiOrbic, the Bioeconomy SFI Research Center of Ireland, manages this program. This research was also supported by the SNSF Ambizione fellowship PZ00-2_233007 awarded to Dr. Viola Vogler-Neuling, the NCCR Bioinspired Materials of the Swiss National Science Foundation (grant number 205603), and the Adolphe Merkle Foundation.

## Author Contributions

Conceptualization: P.K.J.B, L.F., V.V-N. Funding acquisition: P.K.J.B, U.S., V.V.-N. Sample Preparation: P.K.J.B. DNA Extraction: T.D., P.N. Genome Assembly: L.F., Species Identification: C. U. Supervision: U.S., V.V.-N.

Writing– original draft: P.K.J.B, L.F., V.V.-N. Writing – review & editing: all authors.

## References

1. Vogler-Neuling, V. V. et al. Biopolymer Photonics: From Nature to Nanotechnology. Adv. Funct. Mater. 34, 2306528 (2024).

2. Thayer, R. C. & Patel, N. H. A meta-analysis of butterfly structural colors: their color range, distribution and biological production. J. Exp. Biol. 226, jeb245940 (2023).

3. Wilts, B. D., IJbema, N. & Stavenga, D. G. Pigmentary and photonic coloration mechanisms reveal taxonomic relationships of the Cattlehearts (Lepidoptera: Papilionidae: Parides). BMC Evol. Biol. 14, 160 (2014).

4. Zhan, S., Merlin, C., Boore, J. L. & Reppert, S. M. The monarch butterfly genome yields insights into long-distance migration. Cell 147, 1171–1185 (2011).

5. Nowell, R. W. et al. A high-coverage draft genome of the mycalesine butterfly Bicyclus anynana. GigaScience 6, 1–7 (2017).

6. Dasmahapatra, K. K. et al. Butterfly genome reveals promiscuous exchange of mimicry adaptations among species. Nature 487, 94–98 (2012).

7. Mita, K. et al. The genome sequence of silkworm, Bombyx mori. DNA Res. Int. J. Rapid Publ. Rep. Genes Genomes 11, 27–35 (2004).

8. Xia, Q. et al. A draft sequence for the genome of the domesticated silkworm (Bombyx mori). Science 306, 1937–1940 (2004).

9. International Silkworm Genome Consortium. The genome of a lepidopteran model insect, the silkworm Bombyx mori. Insect Biochem. Mol. Biol. 38, 1036–1045 (2008).

10. Miao, X.-X. et al. Simple sequence repeat-based consensus linkage map of Bombyx mori. Proc. Natl. Acad. Sci. U. S. A. 102, 16303–16308 (2005).

11. d’Alençon, E. et al. Extensive synteny conservation of holocentric chromosomes in Lepidoptera despite high rates of local genome rearrangements. Proc. Natl. Acad. Sci. 107, 7680–7685 (2010).

12. Kawahara, A. Y. et al. Phylogenomics reveals the evolutionary timing and pattern of butterflies and moths. Proc. Natl. Acad. Sci. U. S. A. 116, 22657–22663 (2019).

13. Misof, B. et al. Phylogenomics resolves the timing and pattern of insect evolution. Science 346, 763–767 (2014).

14. Ellis, E. A., Storer, C. G. & Kawahara, A. Y. De novo genome assemblies of butterflies. GigaScience 10, giab041 (2021).

15. Wright, C. J. et al. Constraints on chromosome evolution revealed by the 229 chromosome pairs of the Atlas blue butterfly. Curr. Biol. CB 35, 4727-4742.e7 (2025).

16. DNA Extraction from Single Insects. https://assets.ctfassets.net/an68im79xiti/3oGwQ5kl6Uy-CocGgmoWQie/768ae48be4f99b1f984e21e409e801fd/CG000145_SamplePrepDemonstrat-edProtocol_-DNAExtractionSingleInsects.pdf.

17. Miller, S. A., Dykes, D. D. & Polesky, H. F. A simple salting out procedure for extracting DNA from human nucleated cells. Nucleic Acids Res. 16, 1215 (1988).

18. Andrews, S. FastQC: A Quality Control Tool for High Throughput Sequence Data. Babraham Bioinformatics, Babraham Institute (2010).

19. fastp: an ultra-fast all-in-one FASTQ preprocessor | Bioinformatics | Oxford Academic. https://academic.oup.com/bioinformatics/article/34/17/i884/5093234.

20. Cheng, H., Concepcion, G. T., Feng, X., Zhang, H. & Li, H. Haplotype-resolved de novo assembly using phased assembly graphs with hifiasm. Nat. Methods 18, 170–175 (2021).

21. Simão, F. A., Waterhouse, R. M., Ioannidis, P., Kriventseva, E. V. & Zdobnov, E. M. BUSCO: assessing genome assembly and annotation completeness with single-copy orthologs. Bioinformatics 31, 3210–3212 (2015).

22. Rhie, A., Walenz, B. P., Koren, S. & Phillippy, A. M. Merqury: reference-free quality, completeness, and phasing assessment for genome assemblies. Genome Biol. 21, 245 (2020).

23. Ranallo-Benavidez, T. R., Jaron, K. S. & Schatz, M. C. GenomeScope 2.0 and Smudgeplot for reference-free profiling of polyploid genomes. Nat. Commun. 11, 1432 (2020).

24. Brown, M. R., Manuel Gonzalez de La Rosa, P. & Blaxter, M. tidk: a toolkit to rapidly identify telomeric repeats from genomic datasets. Bioinformatics 41, btaf049 (2025).

25. Ou, S. et al. Benchmarking transposable element annotation methods for creation of a streamlined, comprehensive pipeline. Genome Biol. 20, 275 (2019).

26. Edwards, R. J., Dong, C., Park, R. F. & Tobias, P. A. A phased chromosome-level genome and full mitochondrial sequence for the dikaryotic myrtle rust pathogen, Austropuccinia psidii. https://doi.org/10.1101/2022.04.22.489119 (2022) doi:10.1101/2022.04.22.489119.

27. Kurtz, S. et al. Versatile and open software for comparing large genomes. Genome Biol. 5, R12 (2004).

28. Attestad, M. dot. https://github.com/marianattestad/dot

29. Shen, W., Le, S., Li, Y. & Hu, F. SeqKit: A Cross-Platform and Ultrafast Toolkit for FASTA/Q File Manipulation. PLoS ONE 11, e0163962 (2016).

30. De novo genome assemblies of butterflies | GigaScience | Oxford Academic. https://academic.oup.com/gigascience/article/10/6/giab041/6291117.

31. Oh, D.-H. et al. NCBI Orthologs: Public Resource and Scalable Method for Computing High-Precision Orthologs Across Eukaryotic Genomes. J. Mol. Evol. 93, 843–859 (2025).

32. Hu, J. et al. A novel and accelerated method for integrated alignment and variant calling from short and long reads. Front. Bioinforma. 5, (2026).

33. GenomeScope: fast reference-free genome profiling from short reads | Bioinformatics | Oxford Academic. https://academic.oup.com/bioinformatics/article/33/14/2202/3089939.

34. Davey, J. W. et al. Major Improvements to the Heliconius melpomene Genome Assembly Used to Confirm 10 Chromosome Fusion Events in 6 Million Years of Butterfly Evolution. G3 6, 695–708 (2016).

35. Nishikawa, H. et al. A genetic mechanism for female-limited Batesian mimicry in Papilio butterfly. Nat. Genet. 47, 405–409 (2015).

36. Reed, R. D. et al. optix drives the repeated convergent evolution of butterfly wing pattern mimicry. Science 333, 1137–1141 (2011).

37. Zhang, T. et al. Comparison of Long-Read Methods for Sequencing and Assembly of Lepidopteran Pest Genomes. Int. J. Mol. Sci. 24, 649 (2022).

38. Grant, J. R. et al. Proksee: in-depth characterization and visualization of bacterial genomes. Nucleic Acids Res. 51, W484–W492 (2023).

39. Zhang, T. et al. Comparison of Long-Read Methods for Sequencing and Assembly of Lepidopteran Pest Genomes. Int. J. Mol. Sci. 24, 649 (2022).

40. Martin, A. et al. Diversification of complex butterfly wing patterns by repeated regulatory evolution of a Wnt ligand. Proc. Natl. Acad. Sci. U. S. A. 109, 12632–12637 (2012).

41. Muthukrishnan, S., Merzendorfer, H., Arakane, Y. & Kramer, K. Chitin Metabolism in Insects. in Insect Molecular Biology and Biochemistry 193–235 (2012). doi:10.1016/B978-0-12-384747-8.10007-8.

42. Jasrapuria, S., Specht, C. A., Kramer, K. J., Beeman, R. W. & Muthukrishnan, S. Gene families of cuticular proteins analogous to peritrophins (CPAPs) in Tribolium castaneum have diverse functions. PloS One 7, e49844 (2012).

43. Dinwiddie, A. et al. Dynamics of F-actin prefigure the structure of butterfly wing scales. Dev. Biol. 392, 404–418 (2014).

44. Bretscher, A. Deconstructing formin-dependent actin cable assembly. Proc. Natl. Acad. Sci. U. S. A. 110, 18744–18745 (2013).

45. Yu, S., Luo, F. & Jin, L. H. Rab5 and Rab11 maintain hematopoietic homeostasis by restricting multiple signaling pathways in Drosophila. eLife 10, e60870 (2021).

46. Futahashi, R. et al. yellow and ebony are the responsible genes for the larval color mutants of the silkworm Bombyx mori. Genetics 180, 1995–2005 (2008).

47. Reed, R. D. & Nagy, L. M. Evolutionary redeployment of a biosynthetic module: expression of eye pigment genes vermilion, cinnabar, and white in butterfly wing development. Evol. Dev. 7, 301–311 (2005).

48. Liu, S., Wang, M. & Li, X. Overexpression of Tyrosine hydroxylase and Dopa decarboxylase associated with pupal melanization in Spodoptera exigua. Sci. Rep. 5, 11273 (2015).

49. Wang, Q. et al. Tyrosine Hydroxylase and DOPA Decarboxylase Are Associated With Pupal Melanization During Larval-Pupal Transformation in Antheraea pernyi. Front. Physiol. 13, 832730 (2022).

50. Singh, K. S. et al. Genome assembly of Danaus chrysippus and comparison with the Monarch Danaus plexippus. G3 12, jkab449 (2022).

