## Supplementary Information for "Chromosome-level genome assembly of the swallowtail butterfly *Parides eurimedes mylotes* is a valuable resource for studying wing coloration"

Praveen Kumar Jaya Balaji[1,2], Tim Davalan[4], Pamela Nicholson[4], Carmen Rojas Uglade[5], Laurent Falquet[3,6], Viola Vogler-Neuling*[1,2]

1*Adolphe Merkle Institute, University of Fribourg, Chemin des Verdiers 4, 1700 Fribourg, Switzerland.
2 National Center of Competence in Research (NCCR) Bio-Inspired Materials, Chemin des Verdiers 4, 1700 Fribourg, Switzerland
3 Department of Biology, University of Fribourg, Chemin du Musée 10, 1700, Fribourg, Switzerland.
4 Next Generation Sequencing Platform, University of Bern, Bremgartenstrasse 109a, 3012, Bern, Switzerland.
5 Mariposa y Reserva Leonelo Oviedo, Escuela de Biología, Universidad de Costa Rica
6 Swiss Institute of Bioinformatics. 1015 Lausanne, Switzerland

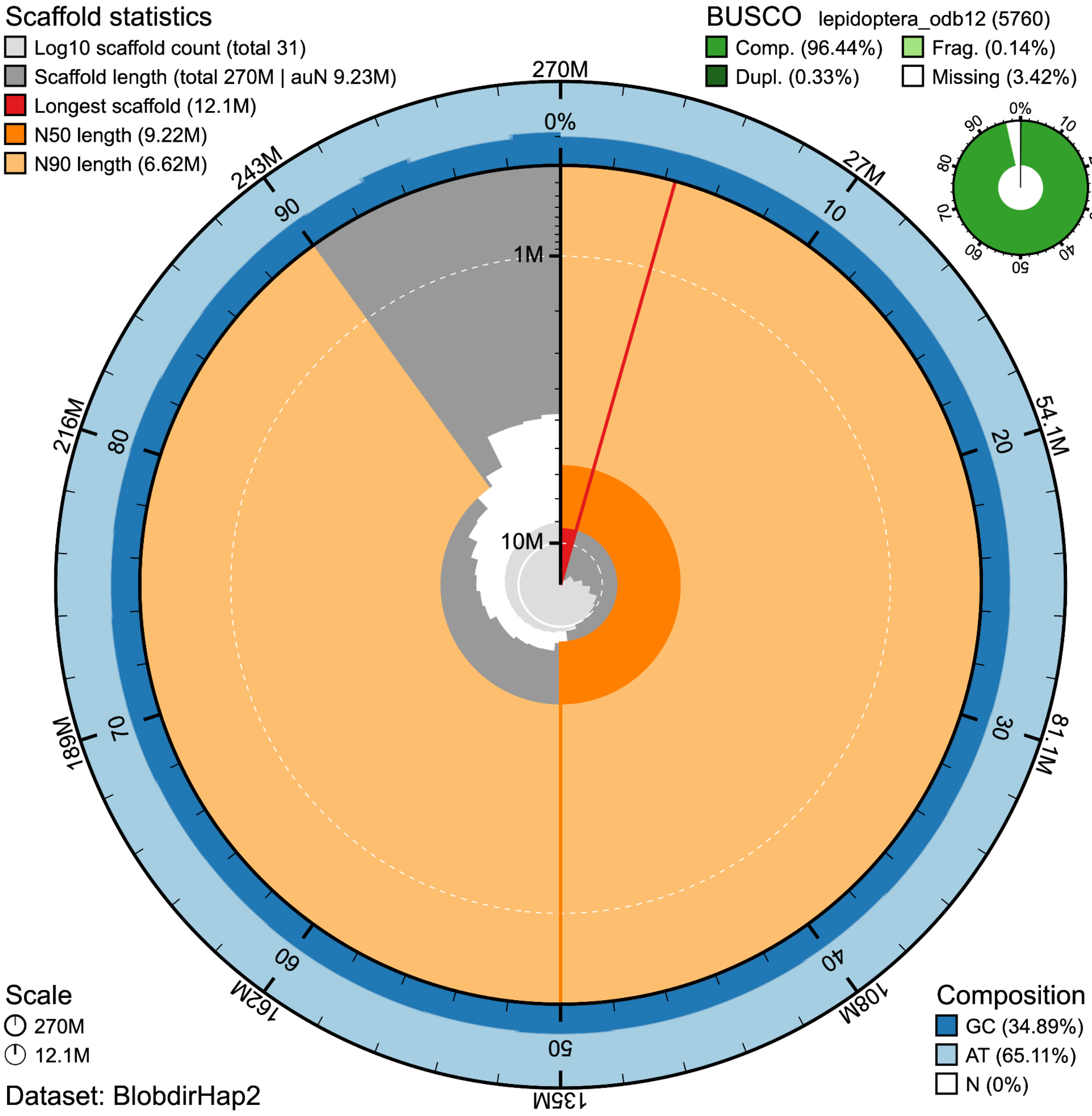


**Suppl. Figure 1:** Snail plots for *Parides eurimedes mylotes* Hap2.

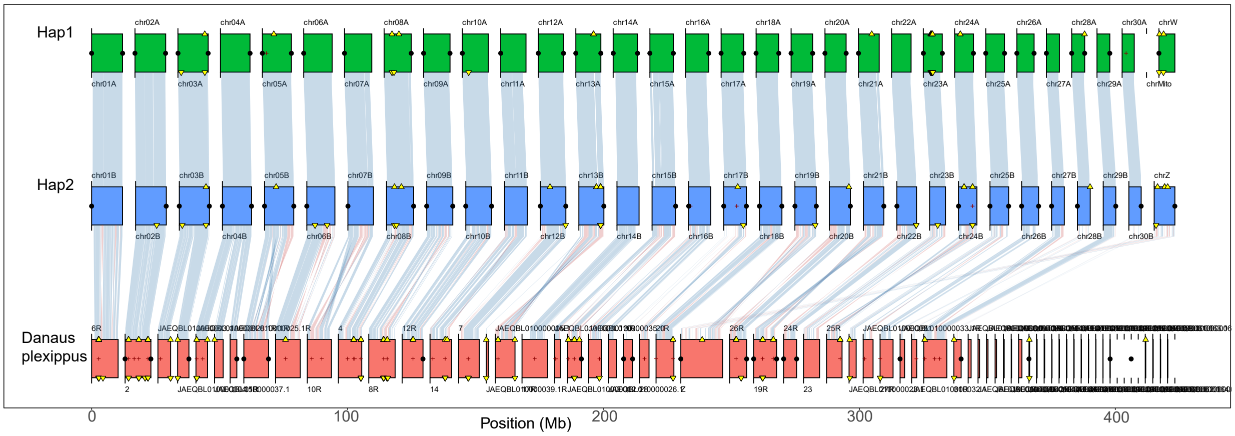


**Suppl. Figure 2:** Synteny plot of the *Danaus plexippus* chromosome (Accession number: GCA_009731565)[50]. Haplotigs 1 (green) and 2 (blue), generated in this study, were tested for synteny against the *Danaus plexippus* reference genome (red).

**Suppl. Table 1:** EDTA output for both haplotigs (Hap1 and Hap2)

| **Repeat Classes Hap1** | | | | |
|---|---|---|---|---|
| ============== | | | | |
| **Total Sequences: 32** | | | | |
| **Total Length: 273940563 bp** | | | | |
| | **Class** | **Count** | **bpMasked** | **%masked** |
| | ===== | ===== | ======== | ======= |
| **LTR** | | -- | -- | -- |
| | Copia | 795 | 489855 | 0.18% |
| | Gypsy | 7772 | 4258293 | 1.55% |
| | unknown | 21417 | 7057170 | 2.58% |
| **TIR** | | -- | -- | -- |
| | CACTA | 27625 | 7189338 | 2.62% |
| | Mutator | 29578 | 5972981 | 2.18% |
| | PIF_Harbinger | 5945 | 1190880 | 0.43% |
| | Tc1_Mariner | 5535 | 949715 | 0.35% |
| | hAT | 7755 | 2033119 | 0.74% |
| **nonTIR** | -- | -- | -- | |
| | helitron | 75941 | 21808126 | 7.96% |
|  | -------------------------------- |  |  |  |
| **total interspersed** | | 182363 | 50949477 | 18.60% |
| **Total** | | 182363 | 50949477 | 18.60% |
| **Repeat Classes Hap2** | | | | |

| | | | | |
|---|---|---|---|---|
| ============== | | | | |
| **Total Sequences: 31** | | | | |
| **Total Length: 270433416 bp** | | | | |
| | **Class** | **Count** | **bpMasked** | **%masked** |
| | ===== | ===== | ======== | ======= |
| **LTR** | | -- | -- | -- |
| | Copia | 341 | 189609 | 0.07% |
| | Gypsy | 7673 | 3993153 | 1.48% |
| | unknown | 15468 | 5160688 | 1.91% |
| **TIR** | | -- | -- | -- |
| | CACTA | 30865 | 8523672 | 3.15% |
| | Mutator | 38229 | 8251752 | 3.05% |
| | PIF_Harbinger | 6563 | 1601596 | 0.59% |
| | Tc1_Mariner | 6036 | 1103126 | 0.41% |
| | hAT | 6828 | 1696506 | 0.63% |
| **nonTIR** | | -- | -- | -- |
| | helitron | 74671 | 20249561 | 7.49% |
|  | --------------------------------- |  |  |  |
| **total interspersed** | | 186674 | 50769663 | 18.77% |
| **Total** | | 186674 | 50769663 | 18.77% |
